# BSReadSim: a versatile and efficient simulator to generate realistic bisulfite sequencing reads

**DOI:** 10.1101/2024.12.24.627620

**Authors:** Wenbin Guo, Matteo Pellegrini

**Affiliations:** Bioinformatics Interdepartmental Program, University of California Los Angeles, Los Angeles, CA, 90095, USA; Department of Molecular, Cell, and Developmental Biology, University of California Los Angeles, Los Angeles, CA, 90095, USA; UCLA-DOE Institute for Genomics and Proteomics, University of California Los Angeles, Los Angeles, CA, 90095, USA

## Abstract

Realistic bisulfite sequencing simulators are crucial for advancing method development in computational epigenetics. However, existing tools often fall short due to oversimplified generative models that fail to capture the complexity of real data. We present BSReadSim, an efficient and versatile simulator that generatesrealistic bisulfite sequencing reads. BSReadSim excels in integrating reference genetic variants and methylation profiles, offering unmatched versatility across multiple sequencing technologies, including WGBS, RRBS, and TBS. By accurately modeling methylation patterns, sampling biases, sequencing errors, and leveraging optimized implementation, BSReadSim efficiently generates realistic synthetic datasets tailored to specific experimental needs while maintaining computational feasibility. By enhancing the realism and flexibility of bisulfite sequencing simulations, BSReadSim supports improved experiment design, method development, and benchmarking of computational tools, ultimately advancing the reliability and rigor of DNA methylation analysis tools.

## Introduction

DNA methylation is a crucial epigenetic modification involving the addition of a methyl group to the fifth carbon of cytosine’s pyrimidine ring. In mammals, this modification predominantly occurs at CpG dinucleotides and plays a critical role in regulating gene expression [1], maintaining genomic stability [2], and underpinning fundamental biological processes such as cellular differentiation [3], development [4], and responses to environmental stimuli [5]. Aberrant DNA methylation has been linked to various diseases [6], establishing it as a central focus in biomedical research for uncovering disease mechanisms and advancing innovative diagnostic and therapeutic strategies. Bisulfite sequencing (BS-seq) is widely recognized as the gold standard for profiling DNA methylation at single-base resolution. In this approach, genomic DNA is treated with sodium bisulfite, where unmethylated cytosines (C) are converted to (U) and subsequently read as thymines (T) during sequencing. In contrast, methylated cytosines (mC) remain unaltered, preserving their sequence identity (C) [7]. This chemical distinction allows for precise differentiation between methylated and unmethylated cytosines, enabling accurate quantification of methylation levels at individual cytosine sites. Despite its unparalleled resolution and accuracy, BS-seq data present significant analytical challenges due to the inherent complexity of DNA methylation dynamics, bisulfite-induced base changes, and technical variability in the sequencing process.

Since the development of bisulfite sequencing, various computational tools have been developed to analyze the bisulfite sequencing data, including specialized read aligners [8–18], SNP-callers [14, 17–23], as well as tools for identifying allelic-specific methylation [22, 24, 25], with nearly every method claiming to achieve the best performance. The pressing need for rigorously benchmarking these tools calls for a reliable simulator to generate realistic bisulfite sequencing data with ground truth. Additionally, a versatile simulator would be invaluable in experimental design. By simulating various scenarios, researchers can determine the best sequencing strategy and optimal sequencing depth needed to obtain accurate methylation measurements, ensuring that experiments are both cost-effective and well-powered. The dual utilities highlight the profound importance of developing a comprehensive bisulfite sequencing simulator to advance both computational tool development and epigenetic research.

Several bisulfite sequencing read simulators have been developed over the years, each tailored to specific technologies with distinct limitations. Among these, Sherman [26], BSBolt [27], and BSSim [28] focus on Whole Genome Bisulfite Sequencing (WGBS). Sherman provides basic simulation functionality but lacks support for genetic variant input and complex methylation profiles. BSBolt allows input of methylation profiles but fails to preserve site-specific methylation levels during simulation. BSSim can support limited genetic variant input through SNP frequency tables, yet it does not incorporate individual genotype data or handle indels. For Reduced Representation Bisulfite Sequencing (RRBS), RRBSsim [29] offers limited support for genetic variants but cannot integrate methylation profiles. Lastly, MethylFASTQ [30] supports both WGBS and Targeted Bisulfite Sequencing (TBS). However, its inability to differentiate between Watson and Crick strands in TBS compromises its utility for targeted sequencing applications. Additionally, like other tools, it does not support genetic variant input or methylation profile simulations. A comprehensive summary of the capabilities and limitations of these simulators is provided in Table 1.

**Table 1:** Summary of existing bisulfite sequencing read simulators.

| Features | Sherman | BSBolt | BSSim | MethylFASTQ | RRBSsim |
| --- | --- | --- | --- | --- | --- |
| Sequencing technology | WGBS | WGBS | WGBS | WGBS/TBS* | RRBS |
| Genetic variant input | No | No | Yes* | No | Yes* |
| Haplotype-aware | No | No | No | No | No |
| Methylation profile input | No | Yes* | No | No | No |
| Site-site dependency | No | No | No | No | No |
| Allelic-specific methylation | No | No | No | No | No |
| Adjustable bisulfite conversion rate | Yes | No | Yes | No | Yes |
| GC-bias/non-uniform coverage | No | No | No | No | No |
| Multi-thread support | No | No | Yes | Yes | No |

(\*) denotes limited support:
- MethylFASTQ cannot distinguish between the Watson and Crick strands for Targeted Bisulfite Sequencing (TBS). - BSSim and RRBSsim only accept SNP input using a frequency table. They cannot faithfully utilize given genotypes, preserve haplotype information, or handle indel variants. - BSBolt randomly picks a value from the methylation reference input for simulation. As a result, for a particular CG site, the simulated data and the reference profile will likely have different methylation levels.

Despite their varying focuses, existing bisulfite sequencing simulators share several critical limitations. First, they fail to fully integrate genetic and epigenetic profiles into the simulated bisulfite sequencing data, constraining their utility in computational tool development, benchmarking, and experimental design. Additionally, these simulators rely on oversimplified generative models, where DNA fragments, base quality scores, and sequencing errors are sampled using uniform probability models. This approach neglects the inherent complexity of bisulfite sequencing data, resulting in less realistic simulations. Furthermore, many of these tools struggle with computational efficiency, and none can simulate data across all three technologies (WGBS, RRBS, and TBS), further limiting their practicality for large-scale applications.

To address these limitations, we propose a novel bisulfite sequencing simulator, BSReadSim, incorporating advanced features such as detailed genetic variant and methylation profile inputs, allele-specific methylation, non-uniform coverage sampling, and quality score and sequencing error modeling. Designed with high-efficiency implementation, the simulator generates realistic bisulfite sequencing data within a practical timeframe. By supporting multiple bisulfite sequencing technologies (WGBS, RRBS, and TBS), BSReadSim provides a robust platform for benchmarking and validating bioinformatics tools under realistic conditions, facilitating experimental design and serving as a valuable resource for computational epigenomic research.

## Results

### Faithful incorporation of reference genetic variants

To evaluate the effectiveness of our bisulfite sequencing reads simulator in integrating genetic variants, we conducted a profile-based simulation using BSReadSim with a customized VCF file aiming at a sequencing depth of 20. After simulation, the synthetic reads were aligned to the reference genome and inspected using the Integrative Genomics Viewer (IGV) [31]. Figure 2 demonstrates the simulator’s ability to faithfully incorporate specified genetic variants into simulated bisulfite sequencing reads.

**Figure 1:**
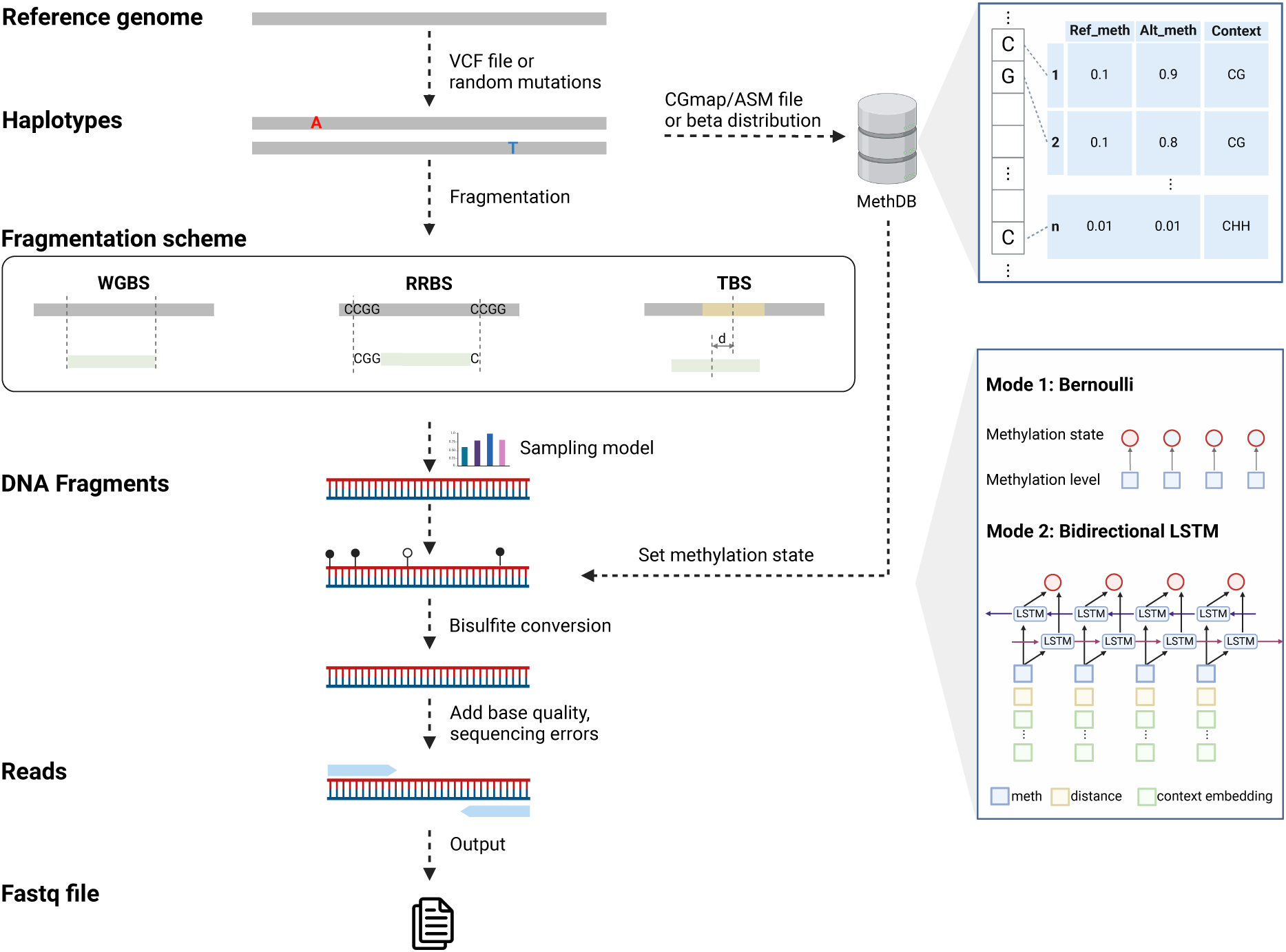
Overview of the Bisulfite Sequencing read Simulation (BSReadSim) Framework. Workflow illustrating the simulation process for bisulfite sequencing data. The process begins with the reference genome, from which haplotypes are generated either through a provided VCF file or by randomly introducing mutations. A methylation database (MethDB) is then constructed, leveraging the methylable bases of the haplotypes and a specified methylation profile (sourced from a CGmap/ASM file or context-specific beta distributions). Subsequently, the haplotypes undergo fragmentation and sampling according to the selected sequencing strategy—WGBS, RRBS, or TBS—to generate DNA fragments. The methylation state of each cytosine within these fragments is determined using a Bernoulli or bidirectional LSTM model. Following the assignment of methylation states, DNA fragments undergo *in silico* bisulfite conversion, read generation, and the addition of base quality scores and sequencing errors to produce realistic bisulfite sequencing reads. Finally, the read data are output in the standardized Fastq file format and ready for downstream analysis.

**Figure 2:**
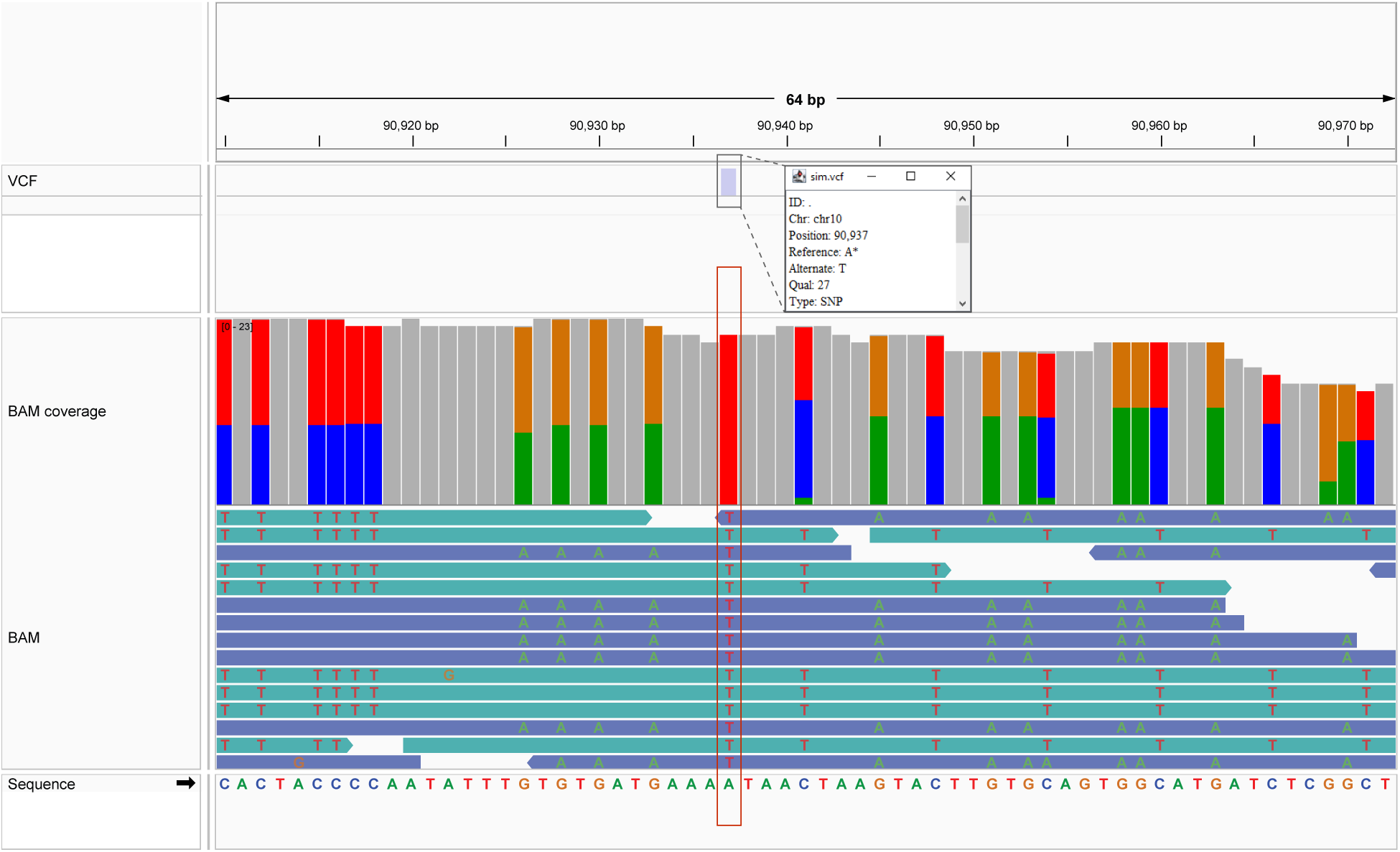
BSReadSim incorporates genetic variants to simulated read data. IGV visualization of read alignment for simulated read data, highlighting the faithful incorporation of predefined genetic variants. The top panel (VCF) displays the predefined VCF file, indicating a homozygous SNP on chromosome 10 at position 90,937, with the reference allele A and the alternate allele T. The middle panel (BAM coverage) illustrates the read coverage at this region, with color-coded bars representing the proportion of reads supporting each base (A: green; C: blue; G: orange; T: red) alongside the sequencing depth distribution. The bottom panel (BAM) shows individual read alignments, where the presence of T alleles at the SNP site is clearly visible, reflecting the homozygous SNP introduced in the simulation and consistent with the input VCF file. The sequence at the bottom of the figure provides the reference sequence context around the SNP. This figure demonstrates an example of the simulator’s capability to faithfully incorporate predefined genetic variants into simulated bisulfite sequencing data.

Specifically, the VCF file contains a homozygous SNP on chromosome 10 at position 90,937, with the reference allele A and the alternate allele T, as shown in the top panel of Figure 2. In the BAM alignment track, all bases at the SNP locus are T, confirming the accurate incorporation of the genetic variant as defined in the VCF file. These results establish our simulator as the first tool to seamlessly integrate predefined genetic variants into bisulfite sequencing reads, offering a reliable platform for advanced epigenomic research. This capability is particularly advantageous for studies that require integrating genetic variants into bisulfite sequencing data, such as developing and benchmarking tools for bisulfite SNP calling, allele-specific methylation, and methylation QTL simulation.

### Accurate preservation of reference methylation profiles

To further evaluate our simulator, we assessed its ability to accurately preserve the reference methylation profiles—a critical feature for generating synthetic data that closely mimics real data and facilitating experimental design. The simulation used a reference genome and a prespecified methylation profile from a CGmap file, targeting at sequencing depth of 20. The generated reads were then aligned to the reference genome, and methylation levels were quantified. Fidelity was evaluated by comparing the designed methylation levels with the estimated levels derived from the simulated data.

We repeat the same simulation procedure for both BSBolt and BSReadSim. The results revealed that BSBolt failed to preserve the reference methylation profile accurately. As shown in Figure 3, the widespread distribution of dots indicates significant discrepancies between the designed and simulated methylation levels. This limitation arises because BSBolt randomly assigns methylation values to CG sites from the reference profile, disregarding their specific genomic location information. In contrast, BSReadSim exhibited much higher fidelity in replicating the reference methylation profile. While minor deviations were observed, primarily due to stochastic variations in sequencing depth, most simulated methylation levels closely aligned with the reference profile, with data points clustering near the diagonal. These results validate BSReadSim as a reliable tool for simulating bisulfite sequencing data, particularly for applications requiring accurate preservation of input methylation profiles.

**Figure 3:**
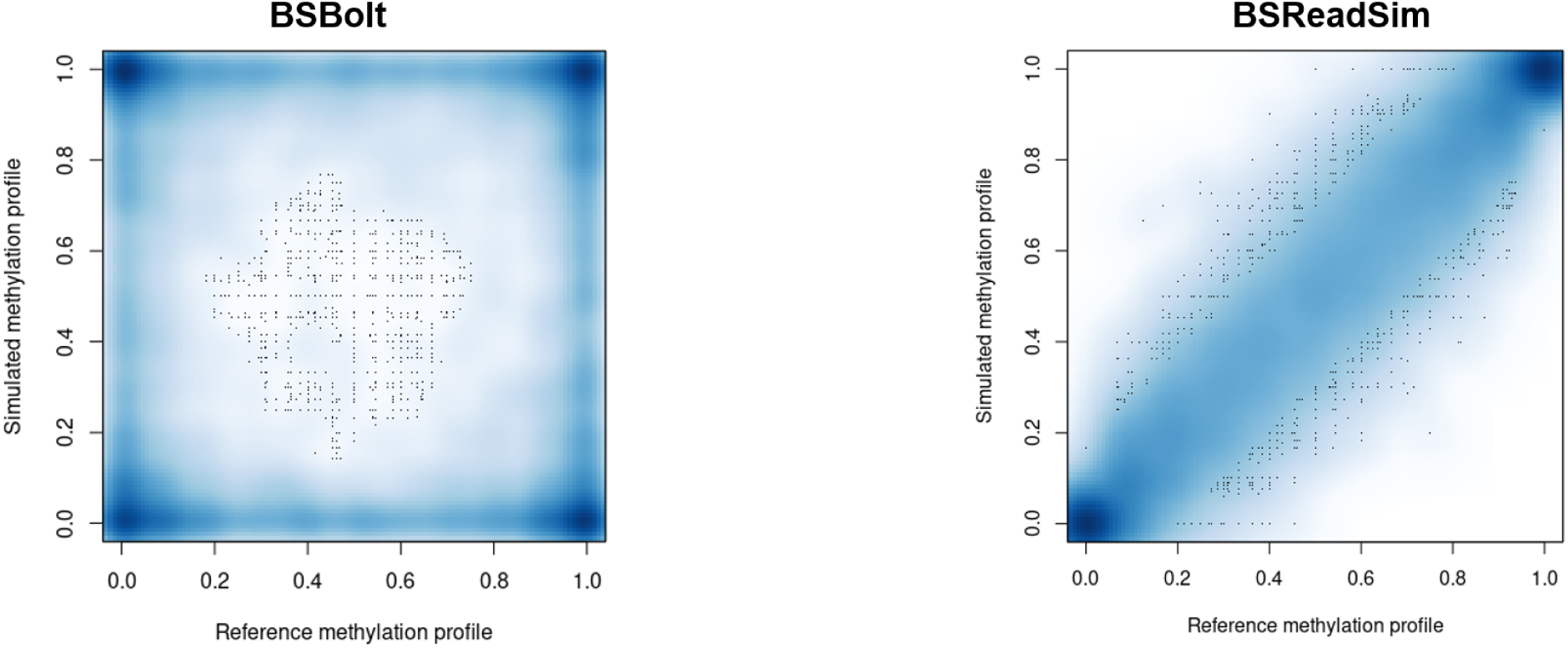
BSReadSim preserves methylation profile in simulated read data. Figure comparing the fidelity of methylation profile preservation between two simulators, BSBolt (left) and BSReadSim (right). Each dot represents a methylable base in the genome, with the x-axis depicting the reference methylation profile and the y-axis showing the methylation profile calculated from the simulated bisulfite sequencing reads. In the BSBolt panel, the widespread distribution of dots indicates a significant loss of location information. Conversely, the BSReadSim panel shows dots closely aligned along the diagonal, demonstrating BSReadSim’s ability to accurately replicate the reference methylation profile and maintain site-specific methylation patterns across the genome. It’s important to note that randomness in sequencing depth can introduce slight variability in the estimated methylation levels, rendering the dots do not perfectly align on the diagonal in the right panel.

### Effective capture of site-site dependency

To evaluate the ability of our BiLSTM-based model to capture site-site dependency, we compared the entropy-distance relationship observed in real data, BiLSTM-simulated data, and Bernoulli-simulated data (Figure 4). Entropy was calculated based on the joint state probabilities of adjacent sites (00, 01, 10, 11), which provides a measure of the methylation concordance in adjacent sites and reflects the site-site dependency. Higher entropy indicates lower concordance between adjacent sites, reflecting weaker site-site dependency. To ensure robust measurement, only site pairs with read counts exceeding 20 were included for analysis.

**Figure 4:**
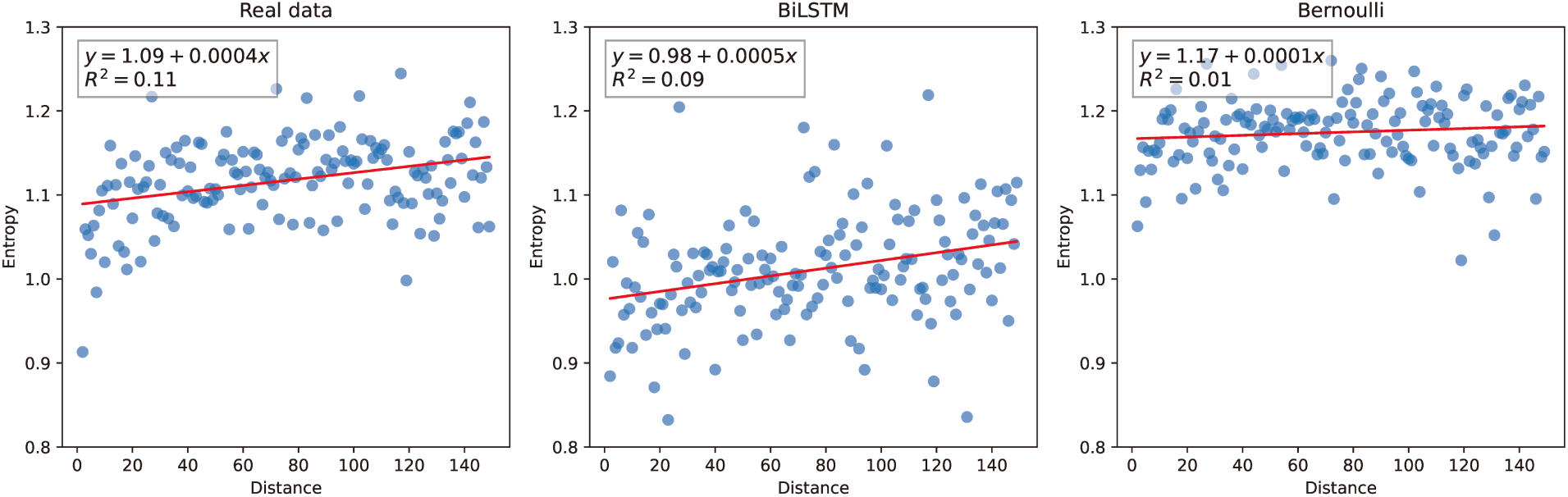
BSReadSim captures of site-site dependency in real data. Comparison of entropy-distance relationships in real data, BiLSTM-simulated data, and Bernoulli-simulated data. The scatterplots depict the entropy of methylation states as a function of the genomic distance between adjacent sites. The red line represents a linear regression fit to the data, with the equation and *R*^2^ value shown in each panel. The left panel shows real data, where a weak but consistent positive correlation is observed (*R*^2^ = 0.11). The middle panel represents data simulated using the BiLSTM model, which closely approximates the real data pattern (*R*^2^ = 0.09), demonstrating its ability to capture site-site dependencies. The right panel shows data generated by the Bernoulli model, which lacks dependency between adjacent sites and exhibits minimal correlation (*R*^2^ = 0.01), highlighting its limitation in reflecting realistic methylation patterns.

In the real data, a weak but significant positive correlation between entropy and distance was observed (*R*^2^ = 0.11), reflecting the gradual weakening of site-site dependencies with increasing distance, consistent with previous finding [32]. The BiLSTM-simulated data closely replicated this trend, showing a comparable positive correlation (*R*^2^ = 0.09), demonstrating the model’s ability to capture realistic dependency structures. However, the entropy in the BiLSTM-simulated data is consistently lower than in the real data, indicating that the real data exhibits greater stochasticity than the model assumes [33]. Future work can consider refining the model to capture the additional variability in the real data not fully captured by the BiLSTM model.

On the other hand, the Bernoulli model assumes that sites are independent, leading to no concordance between adjacent sites. As expected, this results in consistently high entropy that does not vary with distance (*R*^2^ = 0.01). This behavior highlights the limitation of the Bernoulli model in representing the spatial dependencies inherent in real methylation data.

These results emphasize the BiLSTM model’s capability to effectively preserve site-site dependency, making it a valuable tool for generating methylation patterns that reflect biological systems. By capturing both local and long-range dependencies, the BiLSTM-based simulator offers significant advantages over the simpler independent Bernoulli model.

## Discussion

The field of bisulfite sequencing simulation has witnessed the development of several simulators designed to generate synthetic data for various applications. However, existing tools such as Sherman, BSBolt, and MethylFASTQ are limited in their ability to fully integrate genetic and methylation profiles, often producing synthetic data that lacks the complexity of real bisulfite sequencing. These tools also suffer from computational inefficiencies, making them impractical for large-scale studies. To address these limitations, BSReadSim was developed with advanced features, including detailed genetic variant input, allele-specific methylation, and context-aware sequencing error modeling. Results show that BSReadSim can faithfully incorporate reference genetic and methylation profiles while effectively preserve the site-site dependency as observed in real data, providing a robust platform for generating realistic bisulfite sequencing data.

Building on these strengths, BSReadSim can enhance the fidelity of simulations and ensure synthetic data closely mirrors real-world sequencing outputs, making it particularly valuable for benchmarking bioinformatics tools (Figure 5) and designing experiments.

1. **Benchmarking Bisulfite Sequencing Aligners**: Accurate alignment is crucial for down-stream tasks; however, methylation and bisulfite conversion introduce an additional layer of complexity, complicating the alignment process and requiring specialized handling. In the past, a number of aligners have been developed to tackle this challenge. BSReadSim’s ability to generate realistic reads with known fragment origins can provide a rigorous bench-marking framework for bisulfite sequencing aligners, helping identify and refine the most effective alignment tools.
2. **Benchmarking Bisulfite Sequencing SNP Callers**: The identification of SNPs in bisulfite sequencing data is complicated by sequencing errors and bisulfite-induced changes. BSReadSim enables detailed benchmarking of SNP callers by providing synthetic reads that faithfully incorporate genetic variants and provide traceable changes, thereby offering the ground truth necessary for reliable benchmarking of these tools.
3. **Probe Design for Targeted Bisulfite Sequencing and Methylation Arrays**: Designing probes for TBS and methylation arrays requires careful consideration of repetitive elements and potential off-target effects. BSReadSim enables researchers to optimize probe design by simulating TBS data and aligning it back to the reference genome, identifying potential off-target or multi-mapped probes. By providing the feedback loop, BSReadSim can serve to refine the probe sets, thereby improving the reliability and effectiveness of targeted sequencing studies.

**Figure 5:**
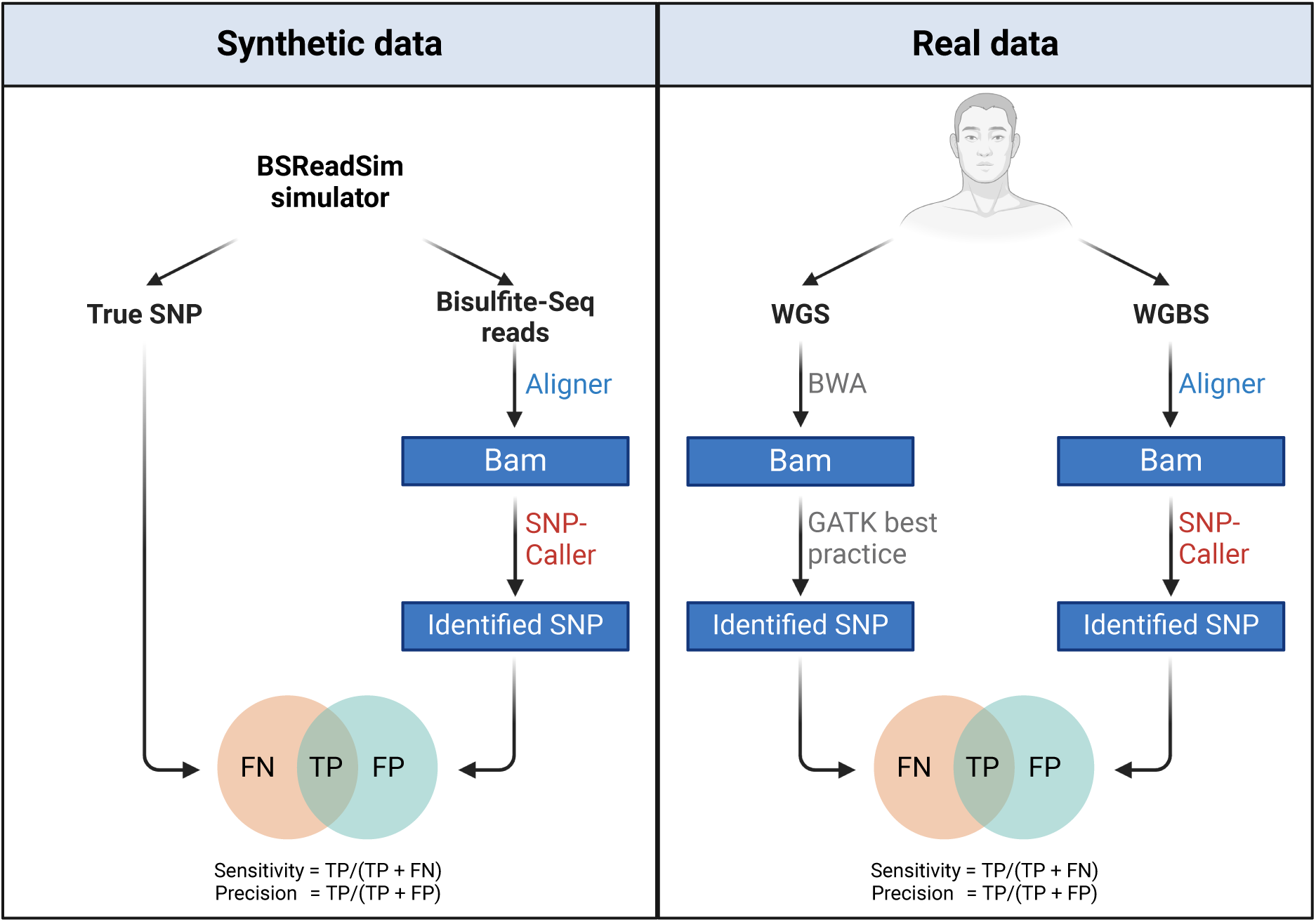
Potential applications of BSReadSim. The left panel illustrates synthetic data generation by BSReadSim with known ground truth, including the true SNPs and the true origin of reads. The synthetic data follows standard bisulfite sequencing read processing steps, including alignment and SNP identification. Each aligner and SNP-caller’s performance can be assessed by comparing the analyzed results to the designed ground truth (benchmarking aligners by comparing the aligned locations to their true origins and benchmarking SNP-callers by comparing identified SNPs to the true SNPs.). The right panel complements this by applying a similar framework to real data from two sequencing modalities (WGS and WGBS), allowing the evaluation of aligner and SNP-calling tools in real-world scenarios. Together, these approaches provide a comprehensive benchmarking framework that integrates both synthetic and real data.

Despite its advancements, several areas remain for improvement and further exploration. BSReadSim currently preserves both genetic and methylation profiles, offering valuable realism for simulating bisulfite sequencing data. However, further testing is needed to assess its unique capability in simulating allele-specific methylation (ASM) [34], which is critical for developing and benchmarking ASM detection tools. Additionally, comprehensive testing and comparisons with existing simulators are necessary to fully evaluate BSReadSim’s computational efficiency and advantages. For site-site dependency modeling, exploring advanced techniques such as Gaussian processes [35] could be further explored to enhance the prediction accuracy. Finally, leveraging BSReadSim to benchmark other bisulfite sequencing tools will be an important step in demonstrating its utility across diverse applications and advancing computational epigenetics research.

In summary, BSReadSim fills a critical gap in bisulfite sequencing by offering a versatile and high-fidelity simulator capable of generating realistic bisulfite sequencing data efficiently. With its unique advantages, BSReadSim supports various applications, including benchmarking alignment and SNP calling tools and optimizing probe design for targeted sequencing experiments. These features highlight its value to the epigenomics research community. Further refinements, including evaluating allele-specific methylation, extensive testing against other simulators, and its use in benchmarking bisulfite sequencing tools, will enhance its impact on computational genetics and epigenomic research.

## Methods

### BSReadSim overview

BSReadSim is designed to generate realistic reads that mimic biological and technical variations observed in real bisulfite sequencing data. The simulator’s workflow, depicted in Figure 1, illustrates the integration of genetic variants, methylation profile, and technical artifacts to produce high-fidelity simulated data. The simulation framework works as follows:

#### Haplotypes generation

The simulator begins with a reference genome sequence from a FASTA file, duplicating each chromosome to represent a diploid organism. Genetic variants, including single nucleotide polymorphisms (SNPs) and short insertions or deletions (indels), can be introduced either by specifying a mutation rate for random mutations or using a pre-defined VCF file. When phased genetic variants are provided in the VCF file, the haplotypes are constructed to accurately reflect the phasing, preserving the true genetic and allelic structure. These haplotypes form the foundation for subsequent simulation steps, including the simulation of allele-specific methylation (ASM).

#### Methylation database construction

Following haplotype generation, a methylation database is created by scanning the methylable bases along the haplotypes and recording their positions and sequence contexts (e.g., CG, CHG, CHH). Methylation levels are assigned to these positions if provided in a CGmap or ASM file. For positions lacking predefined methylation data, context-specific beta distributions, estimated from real methylation profiles, are used to simulate methylation levels, ensuring biologically realistic representation.

#### Fragmentation

The simulator supports three bisulfite sequencing technologies—WGBS, RRBS, and TBS—each employing a tailored fragmentation strategy. DNA fragments are sampled from both haplotypes within a predefined length range (default: 100 to 1000 base pairs). In WGBS, fragmentation sites are randomly distributed across the genome, simulating an unbiased approach. RRBS uses enzyme digestion (e.g., MspI, which recognizes CCGG sites) to concentrate on CpG-rich regions. Thus, DNA fragments are generated between the restriction enzyme sites. For TBS, fragments are centered around provided probe locations, with the exact positions simulated using a Gaussian distribution to account for variability.

#### Methylation pattern assignment

After generating DNA fragments, the simulator assigns methylation patterns to each fragment using the previously constructed methylation database. For each cytosine in a CpG context within the fragment, the methylation level is retrieved from the database, and the methylation state (methylated or unmethylated) is determined using one of two models: an independent Bernoulli model or a bidirectional Long Short-Term Memory (LSTM) network. The Bernoulli model independently assigns each cytosine’s methylation state based on the retrieved methylation level, simulating random methylation patterns. In contrast, the LSTM model incorporates the sequence context and surrounding bases to predict methylation states on the read, leveraging patterns learned from real biological data to simulate biologically realistic methylation states.

#### Bisulfite conversion

After assigning methylation patterns to the DNA fragments, the simulator performs *in silico* bisulfite conversion. In this process, unmethylated cytosines are converted to thymines, while methylated cytosines remain unchanged. To reflect the imperfect nature of bisulfite treatment in real experiments, the simulator applies a fixed conversion success rate, mimicking incomplete conversion. This accounts for scenarios where unmethylated cytosines fail to convert and remain unchanged, ensuring that the final simulated sequences accurately represent both the methylation status and the stochastic nature of bisulfite conversion.

#### Sequencing quality/error assignment

Following bisulfite conversion, the simulator generates a pair of sequencing reads from both ends of the DNA fragments based on the specified read length. Base quality scores are assigned to each base on the reads, either set uniformly across the read or simulated by a Markovian chain using the quality state transition matrix. Sequencing errors are then introduced, either uniformly at a specified error rate or using a quality-specific confusion matrix, ensuring realistic error patterns in the simulated reads.

#### Reads Output and Processing

After sequencing error assignment, the simulator compiles the sequencing reads into standard FASTQ files, including both the nucleotide sequences and their corresponding quality scores. The read name encodes the origin of each read, specifying the chromosome, start, and end positions. Additionally, annotations in the comment line of each read record base changes, such as genetic variants, incomplete bisulfite conversions, or sequencing errors. These detailed annotations provide ground truth, ensuring traceability for each base observation and facilitating downstream benchmarking and analysis.

The generated synthetic reads can be processed and analyzed using the same procedure as the real bisulfite sequencing reads. This may include alignment to a reference genome, methylation and snp calling, or other analyses pertinent to bisulfite sequencing studies. By accurately mimicking the characteristics of real sequencing data, these outputs provide a robust foundation for testing and validating bioinformatics tools and pipelines under controlled conditions. The following sections will detail the modeling component of the simulator to provide a thorough understanding of the simulator’s capabilities and utility, including data sources, modeling details and parameters, as well as implementation specifics.

### DNA fragment sampling model

To accurately replicate the distinct characteristics of different bisulfite sequencing technologies (such as WGBS, RRBS, and TBS), BSReadSim employs DNA fragment generation and sampling processes tailored for each technology. While it also supports the basic uniform sampling approach, as other simulators did, BSReadSim can also offer a profile-based sampling model, where the probability of sampling each DNA fragment is determined by specific fragment features. This allows for more nuanced and realistic data generation.

#### 1. Whole Genome Bisulfite Sequencing (WGBS)

In WGBS, the GC content of a DNA fragment—measured as the proportion of G or C bases—can directly impact its over- or under-representation of fragments in the sequencing output, a phenomenon known as GC bias [36]. The simulator leverages this relationship by using the GC ratio as a primary predictor of sampling probability, expressed as

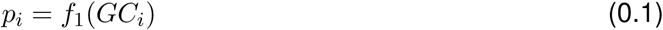

where *GC_i_* represents the GC ratio of the fragment *i*, with sampling probability *p_i_*. We utilized a previously published WGBS dataset from PGP-UK [37] (Sample Accession ID: ERR2359938) to estimate the empirical function *f*_1_. Specifically, the WGBS reads were processed and aligned to the reference genome using BSBolt, which was then divided into 100-base pair windows. For each window, the GC ratio and sequencing depth were calculated. To address variability in sequencing depth, the GC ratio spectrum was divided into 100 bins. Within each bin, the interquartile range (IQR) method was applied to identify and remove regions with extreme sequencing depths, which likely represent alignment artifacts or repetitive elements. The remaining depth values were normalized to scale between 0 and 1 (relative depth), serving as the sampling probability for each region (Figure S1). During simulation, the GC ratio for each fragment is calculated, and rejection sampling is applied based on the corresponding sampling probability. This approach mimics the coverage biases observed in real WGBS data.

### 2. Reduced Representation Bisulfite Sequencing (RRBS)

RRBS targets CpG-rich regions on the genome by utilizing restriction enzymes such as MspI, which cut at specific recognition sites (e.g., CCGG). The simulator replicates this process by first identifying the restriction sites on the haplotypes. It then generates all possible DNA fragments within the predefined fragment length range as candidates based on these restriction sites. The sampling probability for each fragment *i* is modeled as a function of its GC ratio (*GC_i_*), fragment length (*L_i_*), and the number of restriction enzyme sites contained within the fragment (*Count_i_*).

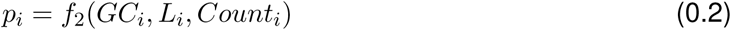

To learn the function *f*_2_, we utilized a previously published RRBS dataset [29]. After processing and aligning the reads to the reference genome, DNA fragments were identified using the read pairs from the RRBS data. For each fragment, sequencing depth, GC ratio, fragment length, and the number of restriction sites were counted. Outliers were removed, and relative depths were calculated using an approach similar to WGBS data. To model the relationship between these features and the observed relative depths, a multivariate spline was fitted (Figure S2), allowing the simulator to estimate the sampling probability for each potential fragment candidate. During the simulation, each fragment was assigned a sampling probability predicted by the model and was subsequently sampled with these probabilities.

#### 3. Targeted Bisulfite Sequencing (TBS)

TBS uses probes to enrich specific genomic regions of interest, with varying capture efficiencies that influence the enrichment of target regions (Figure S3). In BSReadSim, this variability is incorporated by assigning different sampling probabilities to the targeted regions. These probabilities can either be directly provided or empirically estimated from real TBS data to reflect probes’ efficiency or target regions’ accessibility. For empirical estimation, the depth of each probe region is calculated and normalized to generate a relative depth value. During simulation, DNA fragments are sampled from the targeted regions according to their assigned sampling probabilities, ensuring that the simulated data realistically reflects the enrichment and depth variations of real TBS experiments.

### Methylation pattern model

sampled DNA fragments can utilize one of two models to assign methylation states to methylable sites, reflecting varying levels of complexity and realism in simulating methylation patterns.

#### 1. Independent Bernoulli model

The Independent Bernoulli Model is a straightforward approach in which each methylable cytosine site within a DNA fragment is independently assigned a methylation state based on a Bernoulli distribution. The methylation level for site *j*, denoted as *m_j_*, determines the probability of being methylated. The methylation state for site *j* on read *i*, denoted as *y_ij_*, is then determined by:

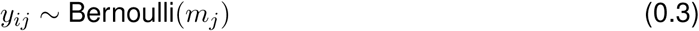

This model is computationally efficient and well-suited for generating baseline methylation patterns. However, it does not consider dependencies between neighboring sites’ methylation states or the influence of genomic context, resulting in less realistic simulated methylation patterns.

#### 2. Bidirectional Long Short-Term Memory (LSTM) model

To account for site-site dependency, we model *Y_i_*, the methylation states of all sites on a read *i*, as being simultaneously sampled from an unknown distribution *g*. This distribution is determined by relevant features, including the methylation levels of sites on the read (*M_i_*), inter-site distances (*D_i_*), and genomic context (*C_i_*), represented as the one-hot embedding of the surrounding sequences. Formally, this can be expressed as:

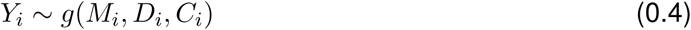

In this work, we utilize a bidirectional LSTM (BiLSTM) model to implicitly learn this function, capturing the intricate relationships among these features. By leveraging its bidirectional architecture, the BiLSTM integrates upstream and downstream sequence and methylation context, allowing it to account for both local and long-range dependencies. This enables the BiLSTM to accurately simulate methylation patterns that reflect the dependencies and variability observed in real biological systems.

To ensure the output methylation states maintain a marginal probability aligned with the predefined methylation levels, we designed a composite loss function combining Binary Cross-Entropy (BCE) loss and Mean Squared Error (MSE) loss. The BCE loss evaluates the accuracy of predicted binary methylation states for each site, while the MSE loss ensures that the averaged prediction state of a read matches the input methylation level. Together, these loss functions guide the BiLSTM in capturing dependencies between adjacent sites while maintaining input methylation levels. This ability to simulate realistic methylation patterns on a read while maintaining methylation level fidelity at individual sites sets our simulator apart from others that lack this capability.

### Sequencing quality and error model

Depending on the user’s needs, the sequencing quality and error on a read can be generated using two approaches: a uniform model and an advanced state transition model. The uniform model assigns a consistent quality score and introduces errors at a constant rate, offering simplicity and computational efficiency. For users requiring greater realism, the advanced model contains the following two parts and captures dependencies across sequential base-calling cycles, providing more realistic simulations.

#### 1. Quality transition matrix

We adopt the same strategy as pIRS [38] and use a quality transition matrix to model quality scores across sequencing cycles. Each element in the matrix represents the probability of transitioning from a quality score in one sequencing cycle to a specific quality score in the subsequent cycle. This approach accounts for the observation that the quality of base calls often depends on the quality of preceding calls, particularly under conditions where the sequencing quality deteriorates along the read. In our simulator, we constructed the quality score transition matrix for read1 and read2, respectively, from the WGBS data, effectively representing the potential difference for the read pairs. (Figure S4)

During the simulation, the quality score for the initial five bases is randomly drawn from the empirical discrete distribution constructed from the real data. For subsequent bases, the simulator uses the quality-transition matrix to determine the following quality score based on the score of the preceding base. This method effectively captures the progressive nature of quality deterioration characteristic of many sequencing platforms, particularly for longer reads.

#### 2. Sequencing error generation

Each quality score has a specific sequencing error profile, with lower quality scores generally indicating higher probabilities of errors and different base errors having different error rates. In our simulator, we empirically derived the base transition matrices for each quality score from real sequencing data. Unlike whole genome sequencing, where error rates can be more directly estimated by comparing the aligned reads to the reference genome, bisulfite sequencing poses additional challenges due to bisulfite conversion, where the observed difference between reads and reference genome can be attributed to either sequencing error or bisulfite conversion. To address this issue, we focus on the overlapped bases of read1 and read2 in paired-end reads. Given the paired reads from the same DNA fragments, any observed discrepancies in the overlapped region must be due to sequencing errors, thus providing a reliable means of estimating errors and minimizing the confounding effects of bisulfite conversion. The estimated error rate profile for each sequencing quality score is presented in Figure S5, effectively capturing the relationship between quality scores and their corresponding error rates.

During simulation, once a quality score is determined for each base, the corresponding base transition matrices are applied to introduce potential sequencing errors using a discrete distribution. This method ensures that the simulated reads realistically reflect the error characteristics observed in actual sequencing experiments.

### Computational optimization strategies

One major bottleneck of bisulfite sequencing read simulation lies in the computational speed. To mitigate this issue, we implemented several computational optimization strategies to ensure efficient processing, enabling the simulation of large datasets within a reasonable timeframe. These strategies are particularly crucial for making the tool accessible and practical for researchers working with high-throughput sequencing data. The following points summarize the key techniques employed:

#### High-efficiency implementation

One key optimization was implementing computational and memory-intensive components of the simulator in C/C++, such as haplotype generation, methylation database construction, and fragment sampling. This lower-level, high-performance language offers better control over memory management and enables efficient data structures, significantly improving computational efficiency. For instance, haplotype construction and the parsing of genetic variants and methylation profiles were implemented using HTSLIB [39], a highly optimized C++ library specifically designed for handling next-generation sequencing data. Additionally, the methylation database was constructed using a customized data structure that utilizes pointers and vectors, ensuring efficient storage and rapid access to site-specific methylation data.

#### Bit encoding and operations

To fully leverage the advantage of C++ and optimize performance, we adopted the bit encoding and operation framework from WGSIM [40], a tool to efficiently simulate whole-genome sequencing reads. By representing each nucleotide (A, C, G, T) as a 2-bit binary value, this encoding reduces the memory footprint by a factor of four compared to traditional byte-based representations. Additionally, bitwise operations—such as AND, OR, XOR, and shifts—are used to perform computations directly on these binary-encoded sequences. These operations are inherently faster than equivalent arithmetic operations by directly manipulating the bits at the hardware level, reducing the time required for tasks such as mutation introduction, fragment generation, and fragment feature extraction. With the compact encoding and the use of fast bitwise operations, BSReadSim not only reduces memory usage but also accelerates computational processes. This dual benefit is particularly important when simulating large genomic datasets with limited computing resources, where both memory efficiency and processing speed are critical.

#### Memory and data access optimization

Efficient memory management was a key focus in our simulator’s design to handle large-scale simulations efficiently and effectively. One of the critical optimizations involved sorting DNA fragments before retrieving their corresponding methylation levels from the methylation database. By sorting fragments, we increased data locality, meaning that related data is accessed sequentially, which significantly reduces cache misses and thus improves processing speed. This approach also exemplifies the trade-off of space for time, as the temporary storage required for sorting is outweighed by the performance gains achieved during data retrieval and processing. On the other hand, the simulator processes genome fragments one chromosome at a time and generates sequencing reads chunk by chunk on the fly, rather than holding all chromosomes and read data in memory simultaneously, reducing the memory footprint and enabling the efficient processing.

#### Algorithmic function optimizations

We identified several frequently used functions in BSReadSim and optimized them for improved performance, such as simulating vectors from Bernoulli and discrete distributions. The Bernoulli distribution is heavily utilized in methylation state assignment and bisulfite conversion; we re-implemented the function by comparing a vector of random numbers to the target probabilities and directly mapping the True/False results to 0/1 states. This optimization achieved a 30-fold speed increase compared to the standard bernoulli.rvs() method for vectors of length 150. The discrete distributions are frequently used for simulating sequencing quality and errors, we optimized the sampling process by generating a uniform random number between 0 and 1 and comparing it against the precomputed cumulative distribution function (CDF) derived from the discrete probabilities. The monotonic nature of the CDF enables efficient identification of the corresponding discrete class using binary search. This optimization reduces computational overhead, significantly accelerating the sampling process, achieving a speed-up of approximately 13.5 times faster than the standard np.random.choice() method.

#### Other optimization endeavors

Beyond the strategies outlined above, we also implemented several additional techniques. The simulator’s scalable and modular design allows efficient handling of datasets ranging from small targeted sequencing to large-scale whole-genome studies, with individual components optimized as needed. Data compression and I/O optimization, such as gzip compression, reduce storage demands and improve data access by processing compressed data directly in memory. Parallel processing further accelerates performance by distributing computational tasks across multiple CPU cores, significantly reducing runtime. These strategies ensure the simulator is both efficient and robust, addressing the demands of high-throughput sequencing simulations.

#### Customizable trade-offs

Recognizing the diverse needs of researchers, we offer users the flexibility to balance computational trade-offs. Users can choose between high-fidelity simulations that prioritize accuracy at the cost of higher resource demands or lower-fidelity simulations optimized for speed. This adaptability allows researchers to tailor simulations to their specific goals and resource availability.

These optimizations significantly enhance the simulator’s performance, making it a valuable tool for the DNA methylome community. By effectively balancing speed and resource efficiency, our simulator provides a robust platform for simulating bisulfite sequencing reads across multiple technologies, experiment design, and the development and testing of bioinformatics methods with high fidelity and realism.

## Code availability

The code is freely available at https://github.com/wbvguo/BSReadSim.git

## Acknowledgements

We thank the public data sources that made this work possible. We also want to thank Harray Zhang, Hongxiang Fu, and Junxi Feng for their valuable input and discussion and the Hoffman2 at UCLA for providing essential computational support. Additionally, we acknowledge ChatGPT for its assistance with debugging and writing. Figure 1 was created using BioRender.com, thus acknowledge it by courtesy.

## Author contributions

W.G. and M.P. conceived the research idea. W.G. implemented the code, performed analysis and wrote the manuscript. M.P. supervised the entire research.

## Supplementary Figures

**Figure S1:**
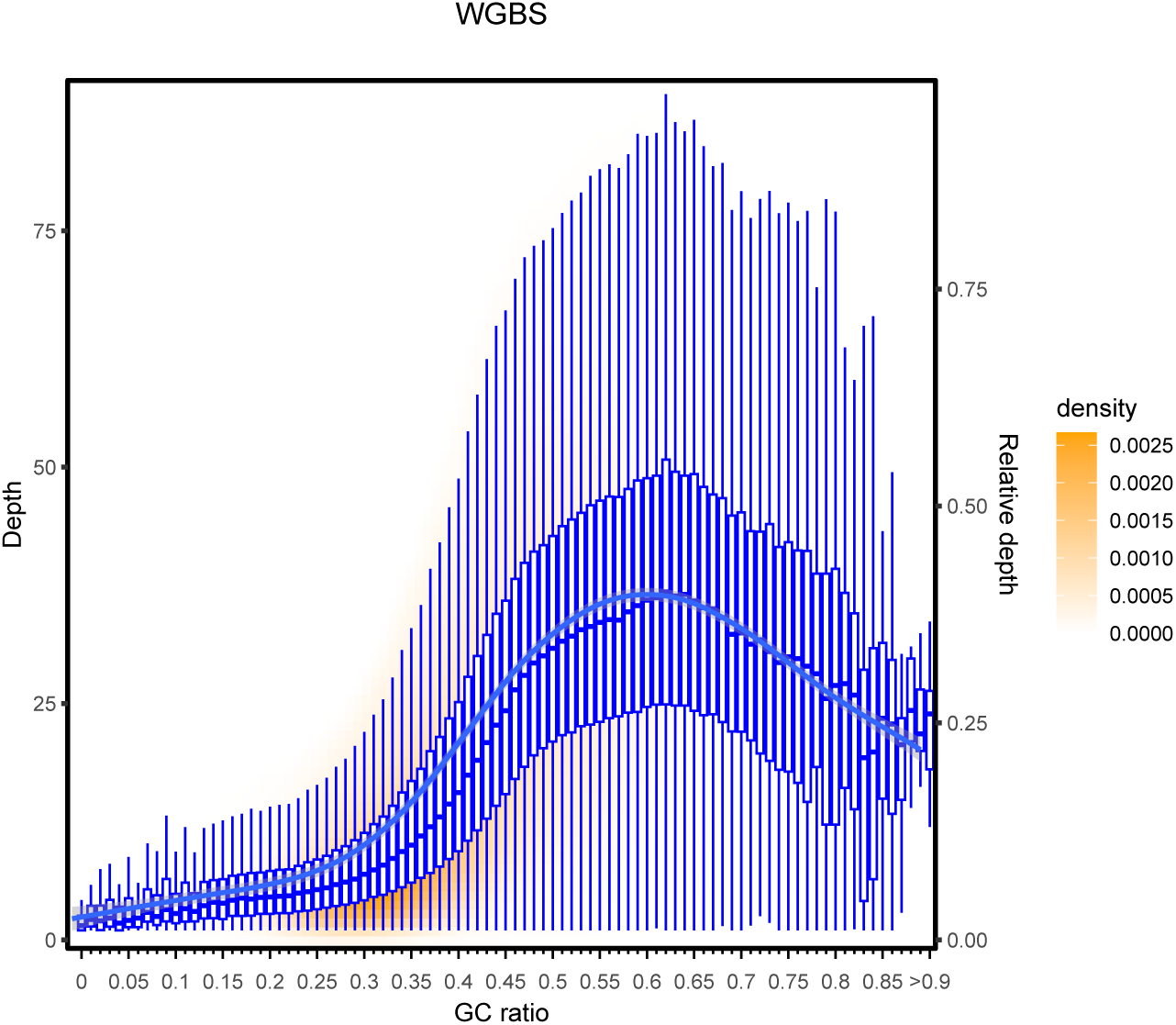
Fragment sampling model for WGBS data. Figure showing the relationship between GC content (x-axis) and relative sequencing depth (right y-axis) in WGBS data. The genome is segmented into 100 bp bins, and the GC ratio and depth are calculated for each bin. Blue boxplots represent the distribution of sequencing depth across different GC ratios, with the central line indicating the median depth, the blue curve representing the local trend, the box representing the interquartile range (IQR), and the whiskers extending to 1.5 times the IQR. The relative depth (right y-axis) is normalized to range between 0 and 1, and the color gradient indicates fragment density (orange for higher density). The figure highlights the GC bias in WGBS, where fragments with intermediate GC content have higher sequencing depth than those with very low or high GC content.

**Figure S2:**
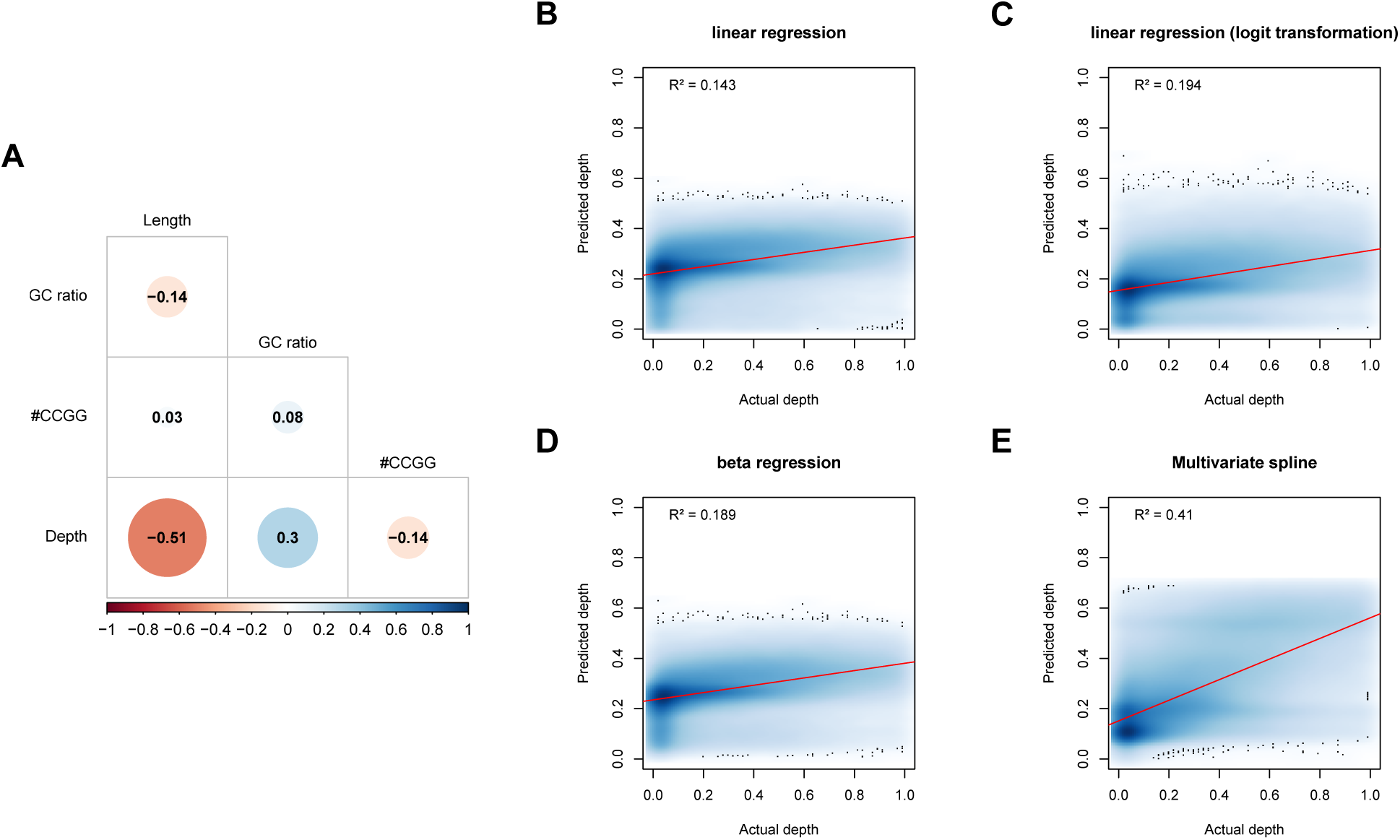
Fragment sampling model for RRBS data. Analysis of factors influencing sampling probability in RRBS data and comparison of predictive models. (A) Correlation matrix showing the relationships between sequencing depth and fragment features (GC ratio, fragment length, restriction site count) in RRBS data. The size and color intensity of the circles indicate the strength and direction of the correlations, with depth showing a negative correlation with fragment length and a moderate positive correlation with GC ratio and the number of restriction sites. (B-E) Comparison of different models for predicting relative depth from the fragment features using linear regression (B), linear regression with logit transformation (C), beta regression (D), and multivariate spline (E). The multivariate spline model (E) shows the best fit, with an *R*^2^ of 0.41, indicating its superior performance in capturing the complex relationship between sequencing depth and fragment features in RRBS data.

**Figure S3:**
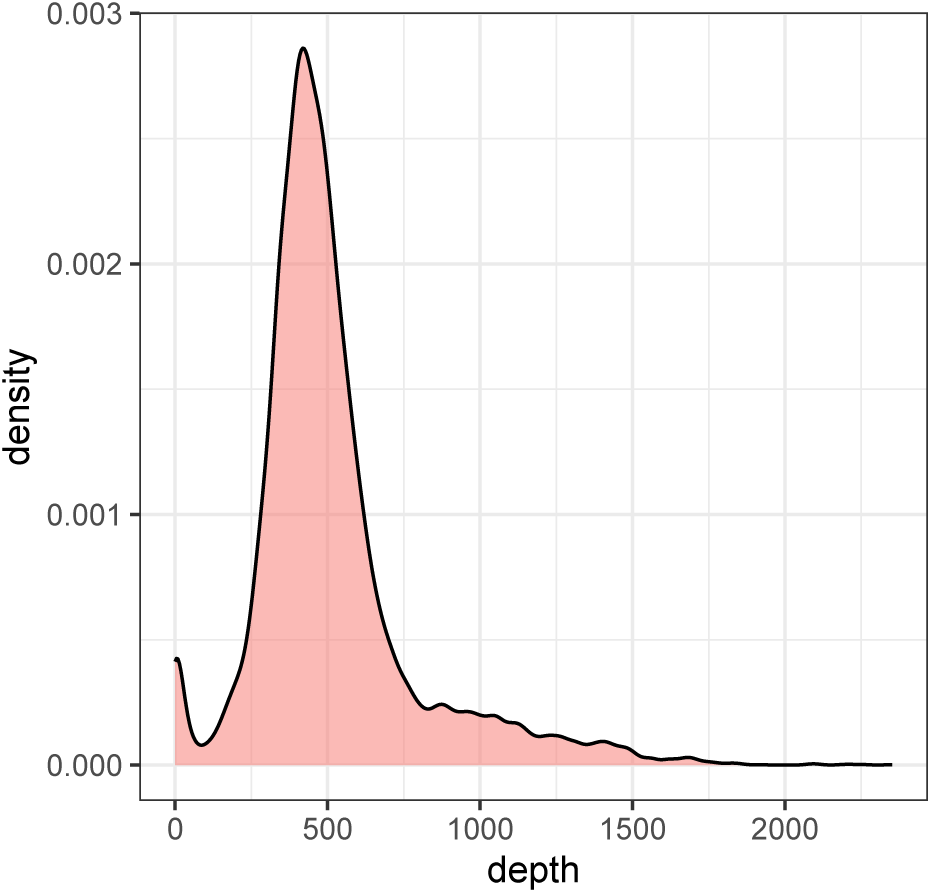
Fragment sampling model for TBS data. Density plot of sequencing depth for targeted regions in Targeted Bisulfite Sequencing (TBS). The plot shows the distribution of sequencing depth across probe-enriched regions, reflecting the variability in capture efficiency of different probes. In the simulation, this variability is modeled by assigning sampling probabilities to targeted regions based on either provided values or empirical estimates from real TBS data. This approach ensures that the simulated reads accurately represent the regional enrichment and depth variations observed in actual TBS data.

**Figure S4:**
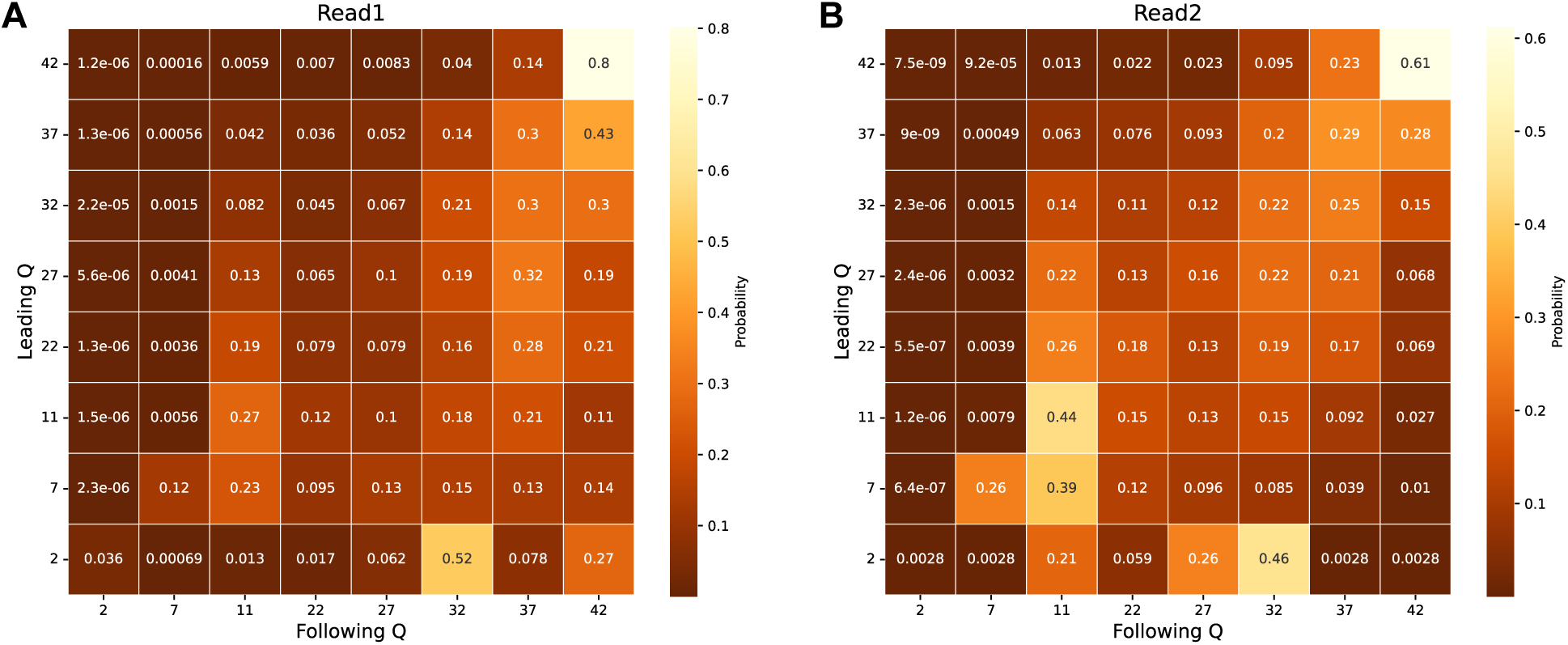
Heatmap of base quality transition probabilities. Quality-transition matrices for Read1 (A) and Read2 (B) in Whole Genome Bisulfite Sequencing (WGBS) data. These matrices represent the probability of transitioning from a preceding quality score (Leading Q) to a subsequent score (Following Q) across sequencing cycles on the read. Each element shows the probability of a quality score change between cycles, with color intensity indicating transition probability. The matrices capture the dependency of sequencing quality on preceding bases and account for quality degradation along read length.

**Figure S5:**
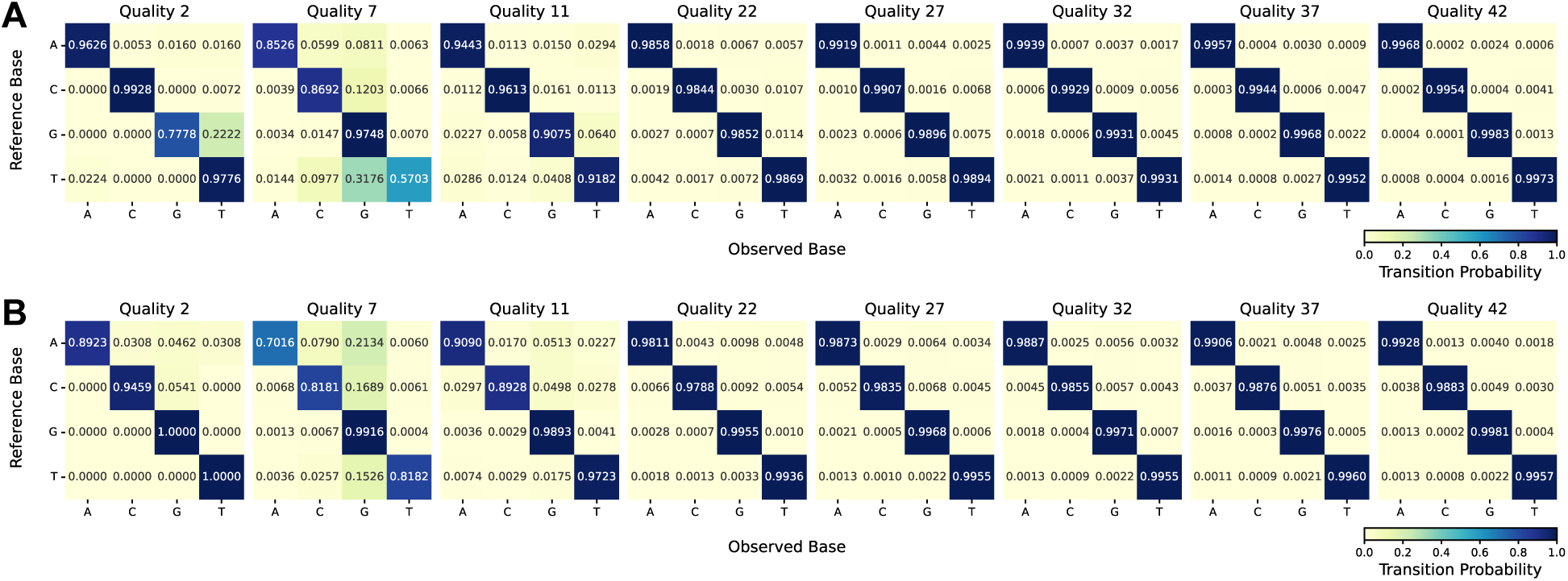
Sequencing error profiles across base quality. Base transition matrices for varying quality scores in bisulfite sequencing data. Panels A and B show the probability of observed bases (A, C, G, T) for given reference bases across different quality scores (2, 7, 11, 22, 27, 32, 37, 42) for Read1 and Read2, respectively. Lower quality scores correspond to higher error probabilities.

## References

[1] Alexandra L Mattei, Nina Bailly, and Alexander Meissner. “DNA methylation: a historical perspective”. In: Trends in Genetics 38.7 (2022), pp. 676–707.

[2] Zachary D. Smith and Alexander Meissner. “DNA Methylation: Roles in Mammalian Development”. In: Nature Reviews Genetics 14.3 (Mar. 2013), pp. 204–220.

[3] Matthias Farlik, et al. “DNA methylation dynamics of human hematopoietic stem cell differentiation”. In: Cell stem cell 19.6 (2016), pp. 808–822.

[4] Maxim V. C. Greenberg and Deborah Bourc’his. “The Diverse Roles of DNA Methylation in Mammalian Development and Disease”. In: Nature Reviews Molecular Cell Biology 20.10 (Oct. 2019), pp. 590–607.

[5] Lisa D Moore, Thuc Le, and Guoping Fan. “DNA methylation and its basic function”. In: Neuropsychopharmacology 38.1 (2013), pp. 23–38.

[6] Zelin Jin and Yun Liu. “DNA methylation in human diseases”. In: Genes & diseases 5.1 (2018), pp. 1–8.

[7] Shawn J Cokus, et al. “Shotgun bisulphite sequencing of the Arabidopsis genome reveals DNA methylation patterning”. In: Nature 452.7184 (2008), pp. 215–219.

[8] Yuanxin Xi and Wei Li. “BSMAP: whole genome bisulfite sequence MAPping program”. In: BMC bioinformatics 10 (2009), pp. 1–9.

[9] Felix Krueger and Simon R Andrews. “Bismark: a flexible aligner and methylation caller for Bisulfite-Seq applications”. In: bioinformatics 27.11 (2011), pp. 1571–1572.

[10] Jing-Quan Lim, et al. “BatMeth: improved mapper for bisulfite sequencing reads on DNA methylation”. In: Genome biology 13 (2012), pp. 1–14.

[11] Weilong Guo, et al. “BS-Seeker2: a versatile aligning pipeline for bisulfite sequencing data”. In: BMC genomics 14 (2013), pp. 1–8.

[12] Brent S Pedersen, et al. “Fast and accurate alignment of long bisulfite-seq reads”. In: arXiv preprint arXiv:1401.1129 (2014).

[13] Elena Y Harris, Rachid Ounit, and Stefano Lonardi. “BRAT-nova: fast and accurate mapping of bisulfite-treated reads”. In: Bioinformatics 32.17 (2016), pp. 2696–2698.

[14] Angelika Merkel, et al. “gemBS: high throughput processing for DNA methylation data from bisulfite sequencing”. In: Bioinformatics 35.5 (2019), pp. 737–742.

[15] Yun Zhang, et al. “Rapid and accurate alignment of nucleotide conversion sequencing reads with HISAT-3N”. In: Genome Research 31.7 (2021), pp. 1290–1295.

[16] Guilherme de Sena Brandine and Andrew D Smith. “Fast and memory-efficient mapping of short bisulfite sequencing reads using a two-letter alphabet”. In: NAR Genomics and Bioinformatics 3.4 (2021), lqab115.

[17] Colin Farrell, et al. “BiSulfite Bolt: A bisulfite sequencing analysis platform”. In: GigaScience 10.5 (2021), giab033.

[18] Wanding Zhou, et al. “BISCUIT: an efficient, standards-compliant tool suite for simultaneous genetic and epigenetic inference in bulk and single-cell studies”. In: Nucleic Acids Research 52.6 (2024), e32–e32.

[19] Yaping Liu, et al. “Bis-SNP: combined DNA methylation and SNP calling for Bisulfite-seq data”. In: Genome biology 13 (2012), pp. 1–14.

[20] Guillermo Barturen, et al. “MethylExtract: high-quality methylation maps and SNV calling from whole genome bisulfite sequencing data”. In: F1000Research 2 (2013).

[21] Shengjie Gao, et al. “BS-SNPer: SNP calling in bisulfite-seq data”. In: Bioinformatics 31.24 (2015), pp. 4006–4008.

[22] Weilong Guo, et al. “CGmapTools improves the precision of heterozygous SNV calls and supports allele-specific methylation detection and visualization in bisulfite-sequencing data”. In: Bioinformatics 34.3 (2018), pp. 381–387.

[23] Adam Nunn, et al. “EpiDiverse Toolkit: a pipeline suite for the analysis of bisulfite sequencing data in ecological plant epigenetics”. In: NAR genomics and bioinformatics 3.4 (2021), lqab106.

[24] Yue Fan, et al. “IMAGE: high-powered detection of genetic effects on DNA methylation using integrated methylation QTL mapping and allele-specific analysis”. In: Genome biology 20 (2019), pp. 1–18.

[25] J Abante, et al. “Detection of haplotype-dependent allele-specific DNA methylation in WGBS data”. In: Nature communications 11.1 (2020), p. 5238.

[26] Babraham Bioinformatics. Sherman. https://github.com/FelixKrueger/Sherman/.

[27] Colin Farrell, et al. “BiSulfite Bolt: A bisulfite sequencing analysis platform”. In: GigaScience 10.5 (2021), giab033.

[28] Qing Xie, et al. “A Bayesian framework to identify methylcytosines from high-throughput bisulfite sequencing data”. In: PLoS Computational Biology 10.9 (2014), e1003853.

[29] Xiwei Sun, et al. “A comprehensive evaluation of alignment software for reduced representation bisulfite sequencing data”. In: Bioinformatics 34.16 (2018), pp. 2715–2723.

[30] Giulia Piaggeschi et al. “MethylFASTQ: a tool simulating bisulfite sequencing data”. In: 2019 27th Euromicro International Conference on Parallel, Distributed and Network-Based Processing (PDP). IEEE. 2019, pp. 334–339.

[31] Helga Thorvaldsdóttir, James T Robinson, and Jill P Mesirov. “Integrative Genomics Viewer (IGV): high-performance genomics data visualization and exploration”. In: Briefings in bioinformatics 14.2 (2013), pp. 178–192.

[32] Ornella Affinito, et al. “Nucleotide distance influences co-methylation between nearby CpG sites”. In: Genomics 112.1 (2020), pp. 144–150.

[33] Andrew E Teschendorff. “On epigenetic stochasticity, entropy and cancer risk”. In: Philosophical Transactions of the Royal Society B 379.1900 (2024), p. 20230054.

[34] Vitor Onuchic, et al. “Allele-specific epigenome maps reveal sequence-dependent stochastic switching at regulatory loci”. In: Science 361.6409 (2018), eaar3146.

[35] Jiawen Chen, et al. “On the identifiability and interpretability of Gaussian process models”. In: Advances in Neural Information Processing Systems 36 (2023), pp. 70267–70278.

[36] Yuval Benjamini and Terence P Speed. “Summarizing and correcting the GC content bias in high-throughput sequencing”. In: Nucleic acids research 40.10 (2012), e72–e72.

[37] Olga Chervova, et al. “The Personal Genome Project-UK, an open access resource of human multi-omics data”. In: Scientific data 6.1 (2019), p. 257.

[38] Xuesong Hu, et al. “pIRS: Profile-based Illumina pair-end reads simulator”. In: Bioinformatics 28.11 (2012), pp. 1533–1535.

[39] James K Bonfield, et al. “HTSlib: C library for reading/writing high-throughput sequencing data”. In: Gigascience 10.2 (2021), giab007.

[40] Heng Li. WGSIM. https://github.com/lh3/wgsim/.

